# Elimination of contamination in the published 16S rRNA gene sequence of *Rothia amarae* type strain J18^T^ during phylogenetic studies of a bacterial isolate from a suspension cell culture of *Arabidopsis thaliana* (Heynh.)

**DOI:** 10.1101/2023.11.08.566172

**Authors:** Sergei Yu. Shchyogolev, Lev A. Dykman, Alexander O. Sokolov, Oleg I. Sokolov, Larisa Yu. Matora

## Abstract

In our phylogenetic studies of a bacterial strain isolated from an *Arabidopsis* suspension culture, we obtained convincing evidence for contamination of the published 16S rRNA gene sequence of *Rothia amarae* type strain J18^T^ (GenBank AY043359.1). Correction of this sequence by deleting the contamination region eliminated contradictions in bioinformatic results that included comparisons of the small-subunit (SSU) rRNA secondary structures. Further, correction of the contaminated sequence yielded sequence identity values for the 16S rRNA genes of the isolate and *R. amarae* type strain J18^T^ (and more than a dozen other *R. amarae* members) above the threshold for species demarcation. The phylogram of the 16S rRNA gene sequences of the type strains closely related to the isolate under study (interspecies sequence identity values, 96–98.7%) united members of the family *Micrococcaceae* (the genera *Rothia, Kocuria* and *Arthrobacter*). A great diversity of habitat conditions was noted for these bacteria, isolated from animals, soil, aqueous media, plant tissues and other sources. This applies, in particular, to *R. amarae* members that are most closely related to the isolate under study by the 16S rRNA gene sequence criterion (sequence identity, near 100%) and belong to four ecotypes: Antarctic, aquatic, soil and endophytic.

## INTRODUCTION

In a suspension culture of *Arabidopsis thaliana* (L.) Heynh (All-Russian Collection of Higher Plant Cell Cultures, Timiryazev Institute of Plant Physiology of the Russian Academy of Sciences, Moscow, Russia), we found the presence of a bacterial microflora (Sokolov *et al*. 2021) that did not have a suppressing effect on the plant. The isolated Gram-positive bacterium was not acid-resistant and had a shape close to spherical (diameter, about 1 μm). The sequencing results for the DNA of the 16S rRNA gene, obtained by the Research and Production Company SYNTOL (SYNTOL 2023), indicate that by this criterion, the isolate is possibly closely related to the species *Rothia amarae*.

*Rothia* is a genus of Gram-positive, aerobic, coccoid or bacillary, nonmotile, non-spore-forming bacteria of the family *Micrococcaceae*, phylum *Actinobacteria* (Austin 2023). To date, 15 *Rothia* species have been identified. *Rothia* are part of the normal microflora of the oral cavity and stomach of humans and animals, but they can cause opportunistic infections of the upper respiratory tract and the gastrointestinal tract in immunocompromised people (Fatahi-Bafghi 2021). The type species, *R. dentocariosa*, was isolated from carious human dentin (Onishi 1949). The complete chromosome-level genome of *R. dentocariosa* type strain ATCC 17931^T^ has reference status for the Human Microbiome Project (ASM16469v2 2023). The GenBank entry for the 16S rRNA gene sequence with a maximum length of 1506 bp, which is 99.87% identical to the same gene from *R. dentocariosa* ATCC 17931^T^, corresponds to the heterotrophic aerobic strain *R. dentocariosa* CV54 (*Rothia dentocariosa* 2023), isolated from saline groundwater in Coimbra, Portugal.

Various *Rothia* isolates have also been recovered from soil, water sources, benthos, rocks, atmosphere, rhizosphere and plant tissues, and other sources (Austin 2023). Some *Rothia* can be used as biofertilizers affecting plant growth and development, as sources of antibiotic peptides, and as degraders of a number of xenobiotics (Austin 2023).

The main object of our studies was to clarify the status of the isolate *R. amarae* sp. nov. (Sokolov *et al*. 2021), hereafter Isolate SG, as a possible true natural symbiont (endophyte; Tadych and White 2019) in a suspension culture of *A. thaliana* (Heynh.), in contrast to the microbial contaminants of cell cultures (Fogh *et al*. 1971). However, in our phylogenetic studies of Isolate SG, we found evidence for contamination of the 16S rRNA gene sequence of *R. amarae* type strain J18^T^ (Fan *et al*. 2002; GenBank AY043359.1), which requires correction.

## MATERIALS AND METHODS

We used the 16S rRNA gene sequence of Isolate SG (Sokolov *et al*. 2021; GenBank OQ702765.1). The strain was deposited in the Collection of Rhizosphere Microorganisms (2023), Institute of Biochemistry and Physiology of Plants and Microorganisms, Russian Academy of Sciences (IBPPM RAS) under number IBPPM 684. We also used the sequences of some known bacterial strains, brought into consideration as a result of the use of the bioinformatic resources mentioned below. The corresponding NCBI accession codes and other characteristics are given in Results and Discussion.

Phylogenetic studies of Isolate SG by the 16S rRNA gene sequence were made with NCBI Standard Nucleotide BLAST (2023). For pairwise comparisons, the option “Align two or more sequences” was used. Multiple sequence alignments (MSAs) were generated with Clustal Omega (2023). Phylogeny.fr (2023) was used to analyze phylogenetic structures. RNA secondary structures for the small subunit of the ribosome (SSU rRNA), encoded by the 16S rRNA gene sequence, were characterized with R2DT (2023), presented by Sweeney *et al*. (2021).

## RESULTS AND DISCUSSION

According to the BLAST results (Standard Nucleotide BLAST 2023), the genera *Rothia, Kocuria* and *Arthrobacter* of the family *Micrococcaceae* were closely related to Isolate SG in the identity *I* of the 16S rRNA gene sequences, provided that *I* > 96%. As stated on the web page (Understanding Results 2023), *I* values of 95–98.6% correspond to strains that should be referred to the same genus. This agrees in principle with the estimates given in Yarza *et al*. (2014), according to which the cut-off threshold for assigning strains to the same genus is *I* = 94.5%.

13 *R. amarae* strains were most closely related to Isolate SG in the 16S rRNA gene sequence identity in the range 99.19–99.91%. However, the *I* of 97.7% between Isolate SG and *R. amarae* type strain J18^T^ (GenBank AY043359.1) was below the intraspecies threshold *I* = 98.65% (Kim *et al*. 2014). This type strain, described in (Fan *et al*. 2002), was isolated from a foul water sewer in Beijing, China, and was described as a Gram-positive bacterium that forms coccoid cells with a diameter of 0.6–0.9 μm, in agreement with the description of Isolate SG in Sokolov *et al*. (2021).

Analysis of the MSA chosen on the basis of the BLAST results (Fig.1) showed that the 16S rRNA gene sequence presented in Fan *et al*. (2002) (GenBank AY043359.1) contained a 20-bp region between 1053 and 1072 bp (CGTGAGATGTCAGCTCGTGT), which most probably resulted from sequencing errors or from contamination.

**Figure 1.**
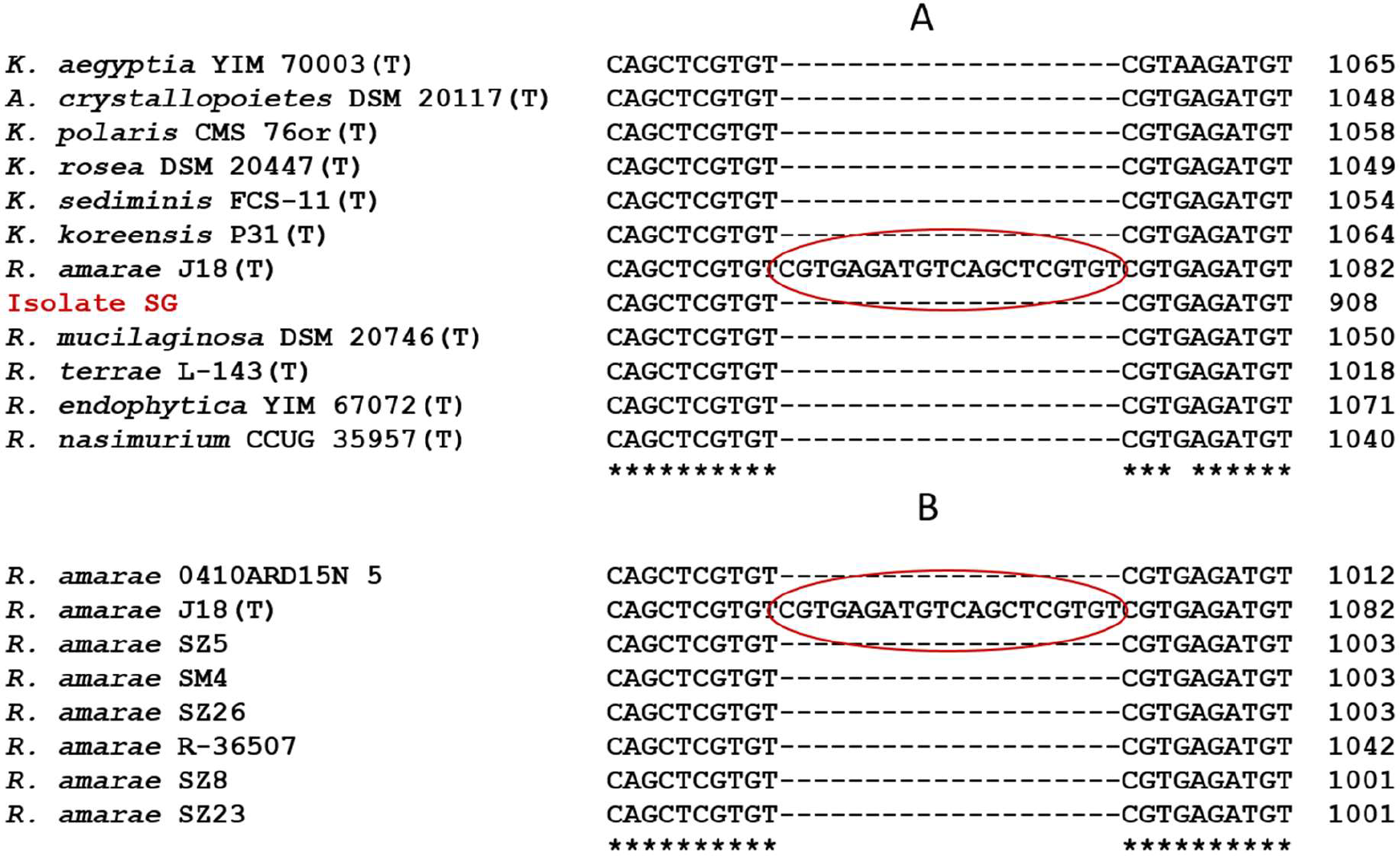
Fragments of the MSAs of the 16S rRNA gene sequences. **(A)** The MSA fragment of *Micrococcaceae* type strains at *I* > 96% with respect to Isolate SG. **(B)** The MSA fragment of *R. amarae* members with a sequence length of at least 1350 bp.

The same conclusion also emerges from the results of comparing R2DT-predicted SSU rRNA secondary structures (R2DT 2023; Sweeney *et al*. 2021) (Fig. 2). In Fig. 2, different colours of letters indicate as follows: black, match to the template; green, modification as compared to the template; red, nucleotide insertion; blue, repositioning relative to the template. With account taken of the genetic sequences, the R2DT program predicts and visualizes secondary (2D) structures of the strains’ SSU rRNA (and other non-coding RNAs) in a standardized position by using appropriate templates, many of which were prepared by experts on the basis of experimental data (Sweeney *et al*. 2021). The templates are selected either automatically (by default) or manually from a special list of templates provided. The results of Sweeney *et al*. (2021), based on a large factual material, show close correspondence between R2DT predictions and selected templates (more than 90% of all nucleotides within the sequences were in the positions corresponding to those of the templates), despite the considerable phylogenetic distances between the templates and the analyzed RNA sequences.

**Figure 2.**
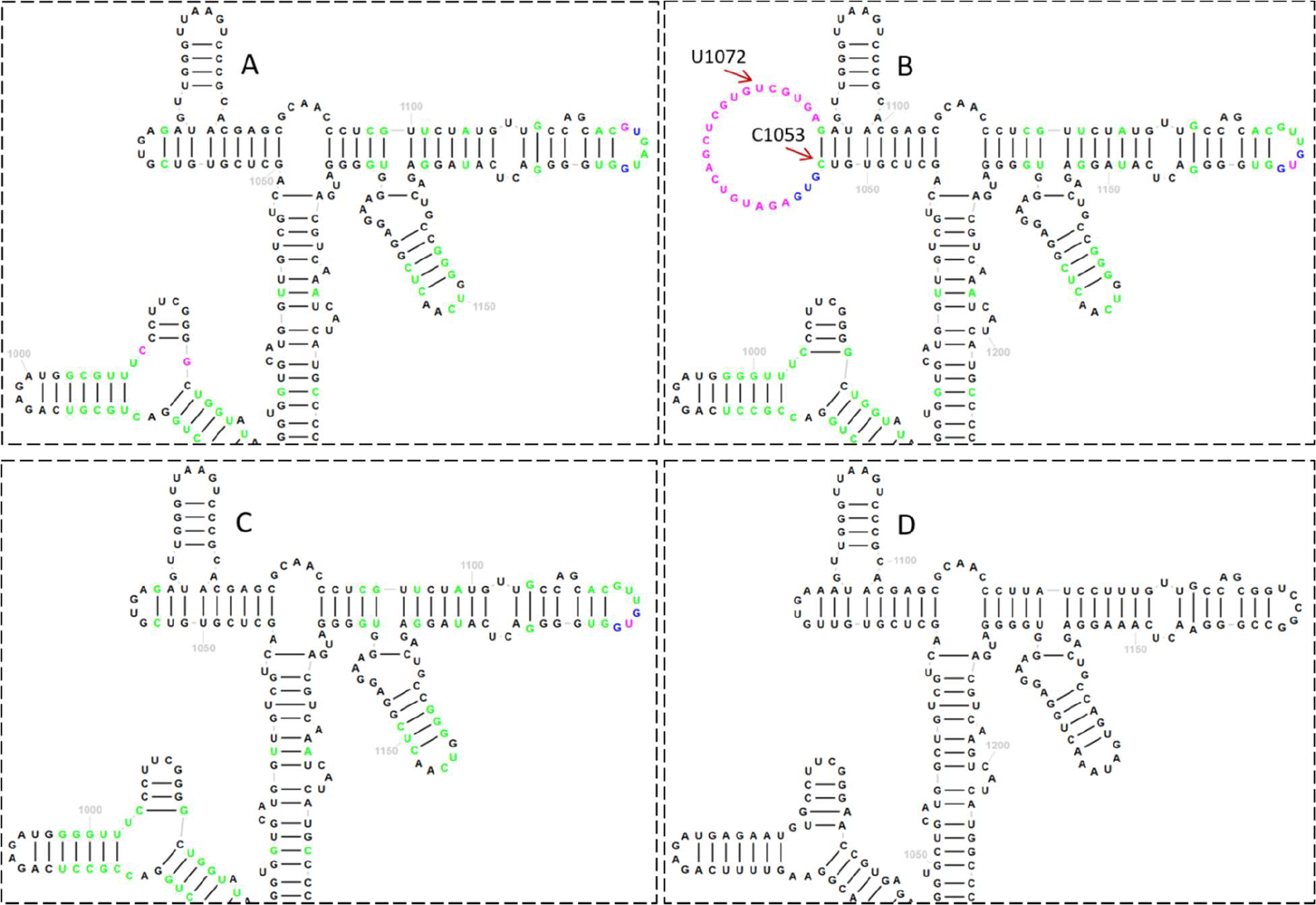
Fragments of the SSU rRNA 2D structures, as predicted by R2DT (2023). **(A)** For the 16S rRNA gene sequence of *R. amarae* J18^T^ with the deleted 20-bp region, highlighted in Fig. 1. **(B)** For the original 16S rRNA gene sequence of *R. amarae* J18^T^. **(C)** For the 16S rRNA gene sequence of *R. dentocariosa* ATCC 17931^T^, a member of the type species of *Rothia*. **(D)** For the 16S rRNA gene sequence of *Escherichia coli* K12, whose 2D SSU rRNA secondary structure was used by the program as a template. Explanations are in the text.

This can also be seen in our Fig. 2 results. The overall structure of the SSU rRNA fragments (and 2D structures in general; not shown here) is almost identical for members of the genera *Rothia* (Gram-positive bacteria, class *Actinomycetes*; Fig. 2A–C) and *Escherichia* (Gram-negative bacteria, class *γ-Proteobacteria*; Fig. 2D). Also preserved are the nucleotide modifications (green colour) in *R. amarae* J18^T^ (Fig. 2A,B) and *R. dentocariosa* ATCC 17931^T^, as compared to the template (Fig. 2C). Against this general background stands out a loop that corresponds to a putative 20-bp-long contaminant (Fig. 1), marked in red by R2DT in Fig. 2B and identified as a nucleotide insertion. This loop, which is absent in Fig. 2A,C,D, is also absent in the SSU rRNA secondary structures for other *Micrococcaceae* studied in our work (data not shown). A perturbation of the SSU rRNA secondary structure for *R. amarae* J18^T^, associated with the C1053–U1072 loop (Fig. 2B) and undetected in other strains, could strongly affect ribosome functioning and, apparently, should be regarded as scarcely probable.

Thus, the results in Figs. 1 and 2 give grounds to edit the 16S rRNA gene sequence of *R. amarae* J18^T^ (GenBank AY043359.1) by deleting the 20-bp-long region identified in Fig. 1. Note that after the deletion of the putative contamination site from the 16S rRNA gene sequence of *R. amarae* J18^T^, the 16S rRNA gene sequence identity between *R. amarae* J18^T^ and Isolate SG (99.73%) exceeds the 98.65% threshold for species demarcation (Kim *et al*. 2014). This is the basis for assigning Isolate SG by the 16S rRNA gene sequence criterion to the species *Rothia amarae* and giving it the strain name *Rothia amarae* SG (GenBank OQ702765.1). In all further bioinformatic computations, we used the edited 16S rRNA gene sequence of *R. amarae* strain J18^T^.

The bacterial 16S rRNA gene sequence contains nine markedly hypervariable regions (V1–V9) (Gray *et al*. 1984; Chakravorty *et al*. 2007; Yang *et al*. 2016), and this crucially determines the potential of 16S rRNA gene sequence as a popular phylogenetic marker. Different sites differ in the degree of conservation, which ensures their phylogenetic significance for different taxonomic categories—from phylum and order to genus and species (Gray *et al*. 1984; Chakravorty *et al*. 2007; Yang *et al*. 2016). According to the results of Chakravorty *et al*. (2007), regions V2, V3 and V6 are the most informative in phylogenetic tests at the genus and species levels.

Consequently, in phylogenetic studies that are aimed at species identification of isolates (or metagenomic objects) and use 16S rRNA gene sequences whose length often differs considerably from the maximum length (about 1500 bp), the effectiveness depends on the presence in them of a sufficiently representative set of hypervariable specific regions, primarily those noted above.

To clarify this with respect to the 16S rRNA gene sequence of Isolate SG (GenBank OQ702765.1), we used a comparison of the SSU rRNA secondary structure for *R. amarae* J18^T^ with the corresponding reference structure of *E. coli* K12 (16S ribosomal RNA 2023), with a known distribution in it of elements corresponding to regions V1–V9 (Fig. 3). We took into account the results of pairwise alignment of the 16S rRNA gene sequences of *R. amarae* J18^T^ and Isolate SG, which had been obtained with Standard Nucleotide BLAST (2023) and had shown their identity *I* = 99.73% (see above).

**Figure 3.**
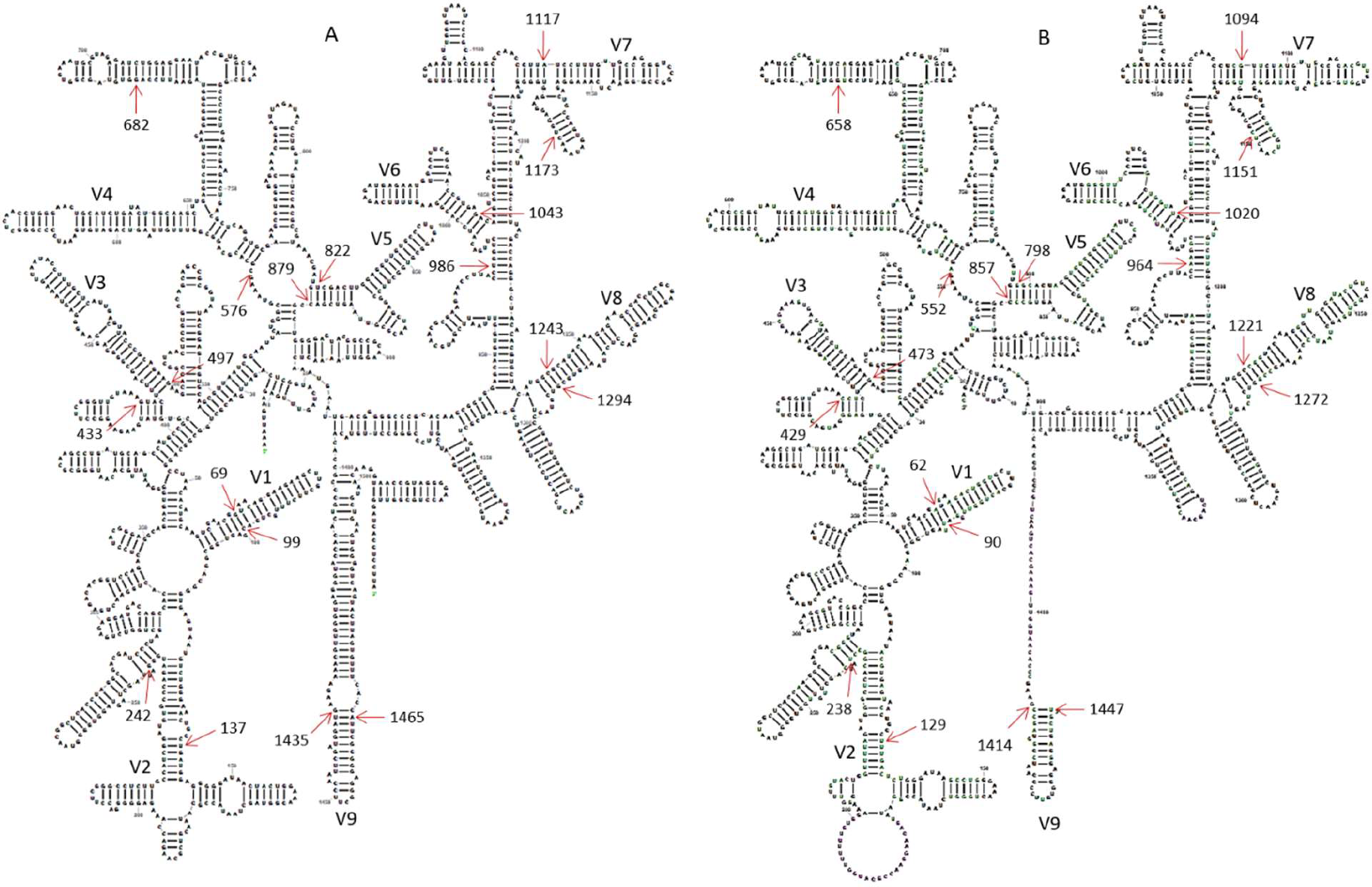
SSU rRNA 2D structures predicted with R2DT (2023). **(A)** For the 16S rRNA gene sequence of *E. coli* K12 (16S ribosomal RNA 2023). **(B)** For the edited 16S rRNA gene sequence of *R. amarae* J18^T^. Numbers with arrows mark the boundaries of hypervariable regions V1–V9. Explanations are in the text.

The numbering of the nucleotide residues in regions V1–V9 for *E. coli* K12 in Fig. 3A corresponds to the results of Chakravorty *et al*. (2007). The high conservation of the secondary structures of prokaryotic SSU rRNAs across long evolutionary distances (Sweeney *et al*. 2021), which is shown, in particular, in Fig. 3, allows us to project the boundaries of regions V1–V9, numbered in the 16S rRNA gene sequence of *E. coli* K12 (Fig. 3A), to similar regions numbered in the edited 16S rRNA gene sequence of *R. amarae* type strain J18^T^ (Fig. 3B): V1, 62–90; V2, 129–238; V3, 429–473; V4, 552–658; V5, 798–857; V6, 964–1020; V7, 1094–1151; V8, 1221–1272; and V9, 1414–1447.

Note that the numbering of the boundaries of regions V1–V9 in the 16S rRNA gene sequence of *R. amarae* J18^T^, obtained from analysis of the SSU rRNA secondary structures (Fig. 3), is confirmed by the MSA results with 16S rRNAgene sequences of *E. coli* K12, *R. amarae* J18^T^ and Isolate SG, obtained with Clustal Omega (2023) (data not shown).

By aligning the 16S rRNA gene sequences of *R. amarae* J18^T^ and Isolate SG with Standard Nucleotide BLAST (2023), we obtained projections of the hypervariable regions onto the 16S rRNA gene sequence of Isolate SG. The results in Fig. 4 show wide coverage of hypervariable regions by the 16S rRNA gene sequence of Isolate SG, except for the relatively short terminal regions V1 and V9, which are inessential for bacterial species identification (Chakravorty *et al*. 2007; Yang *et al*. 2016). This shows the feasibility of using this sequence in the phylogenetic studies of Isolate SG presented in this work.

**Figure 4.**
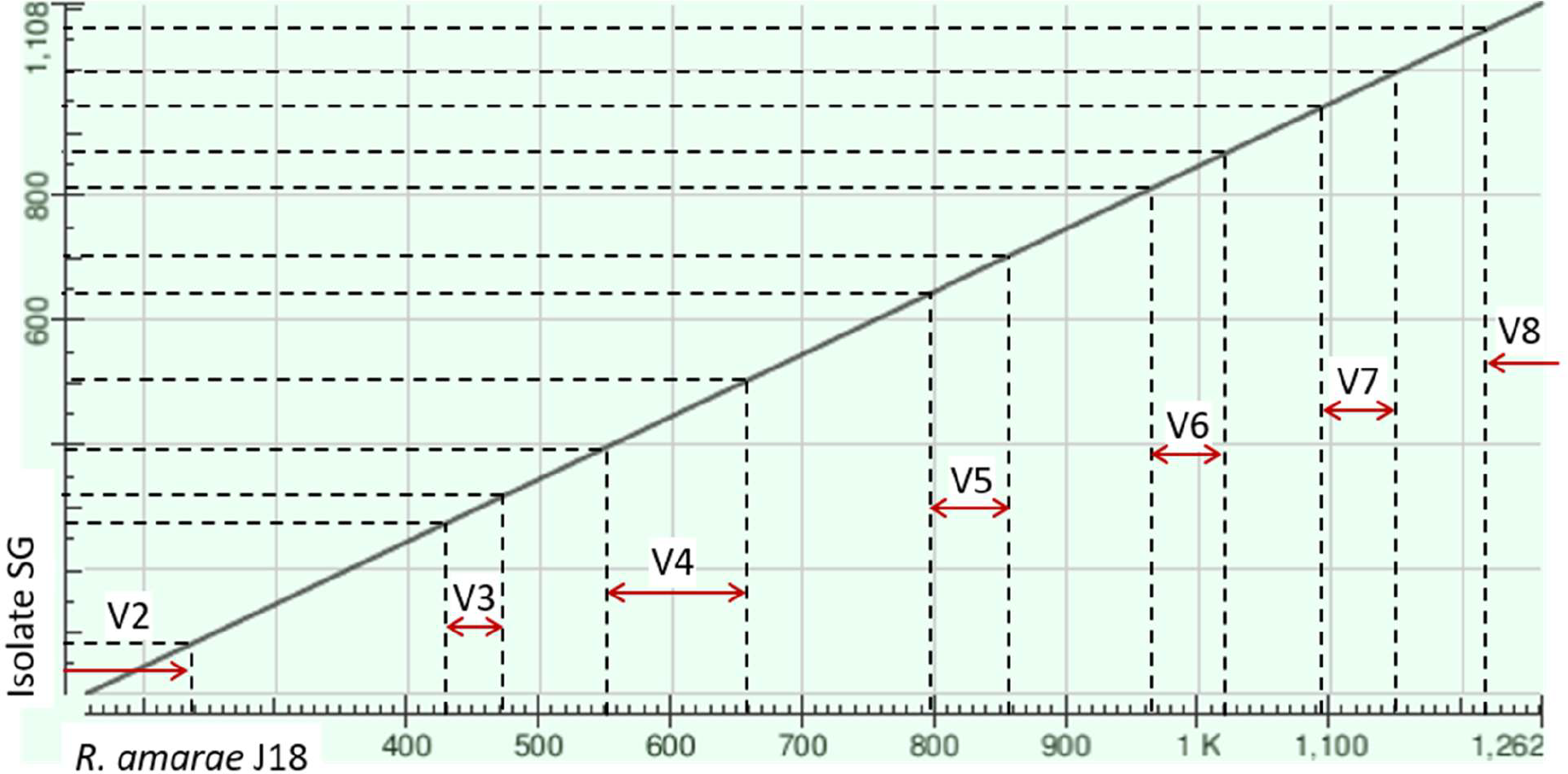
Identification of hypervariable regions in the 16S rRNA gene sequence of Isolate SG by Standard Nucleotide BLAST (2023). The results of their identification in the 16S rRNA gene sequence of *R. amarae* J18^T^ with the aid of the SSU rRNA secondary structures of the strains (Figure 3) were used. Sequence identity *I* = 99.73%.

The use of BLAST for the 16S rRNA gene sequence of Isolate SG led us to identify 14 *R. amarae* strains with an identity of 99% < *I* < 100% above the *I* = 98.65% threshold for joining strains within a species (Kim *et al*. 2014) (Table 1). However, these strains differ substantially in that they occupy 10 ecological niches extending from sewage and ocean water to plant tissues, soil and rocks in Antarctica. Obviously, the existence of bacteria in such contrasting habitats should be ensured by the corresponding sets of phenotypes encoded in their genomes, including the essential role of horizontal gene transfer (Wiedenbeck and Cohan 2011; Koonin 2012). The latter cannot be accounted for in phylogenetic studies using markers from the core component of the pangenome—’housekeeping’ genes (Koonin 2012)—including 16S rRNA gene sequences. Following the considerations given in Wiedenbeck and Cohan (2011), Cohan (2002) and Cohan (2017), we can state the existence of four *R. amarae* ecotypes: Antarctic, aquatic, soil and endophytic.

**Table 1.** Results of a BLAST pairwise alignment of the 16S rRNA gene sequences of Isolate SG with those of *R. amarae* strains differing in their habitat.

| No. | Strain | GenBank accession | Length (bp) | I (%) | Isolation source |
| --- | --- | --- | --- | --- | --- |
| 1 | HMB110 | MG905369.1 | 1419 | 99.91 | Sponge in the Yellow Sea, China |
| 2 | SEFSK10 | KU605695.1 | 1322 | 99.91 | Sweet pepper, Budapest, Hungary |
| 3 | SZ23 | LT600551.1 | 1369 | 99.82 | Seawater in the southern Indian Ocean |
| 4 | SZ8 | LT600537.1 | 1381 | 99.82 |  |
| 5 | SM4 | LT600586.1 | 1371 | 99.73 |  |
| 6 | SZ5 | LT600534.1 | 1372 | 99.73 |  |
| 7 | SZ26 | LT600554.1 | 1372 | 99.64 |  |
| 8 | Bil-LT4_1 | MZ469307.1 | 1400 | 99.82 | Bilberry leaves of the Baltic-Nordic region, Lithuania |
| 9 | KJZ-9 | CP061538.1 | 1530 | 99.82 | Soil dirt, Shanghai, China |
| 10 | J18<sup>T*</sup> | AY043359.1 | 1447 | 99.73 | Foul water sewer in Beijing, China |
| 11 | R-36507 | FR682692.1 | 1482 | 99.73 | Soil sample at the Princess Elisabeth Station, East Antarctica |
| 12 | SMV513-R2_L30 | OP364104.1 | 1362 | 99.73 | Soil sample, Milan, Italy |
| 13 | P00107Karwar | MG800697.1 | 1352 | 99.64 | Marine sediments at an open sea cage farm on the west coast of India |
| 14 | 0410ARD15N_5 | FR848426.1 | 1426 | 99.19 | Air from a show cave in Spain |

\* Edited sequence

To evaluate the systematic position of Isolate SG in terms of 16S rRNA gene sequences at evolutionary distances corresponding to interspecies identity values of 96% < *I* < 98.65% (Yarza *et al*. 2014; Kim *et al*. 2014), we chose 11 type strains (Table 2) on the basis of the BLAST results. Fig. 5 is a phylogram of 16S rRNA gene sequences, obtained with Phylogeny.fr (2023), which characterizes the taxonomic environment of Isolate SG under the above conditions. To the ecological niches of the above *R. amarae* strains, which are most closely related to Isolate SG by 16S rRNA gene sequences (Table 1), are added various habitats characteristic of micrococci. These habitats include animals, soil, sediments of water bodies and fermented seafood.

**Figure 5.**
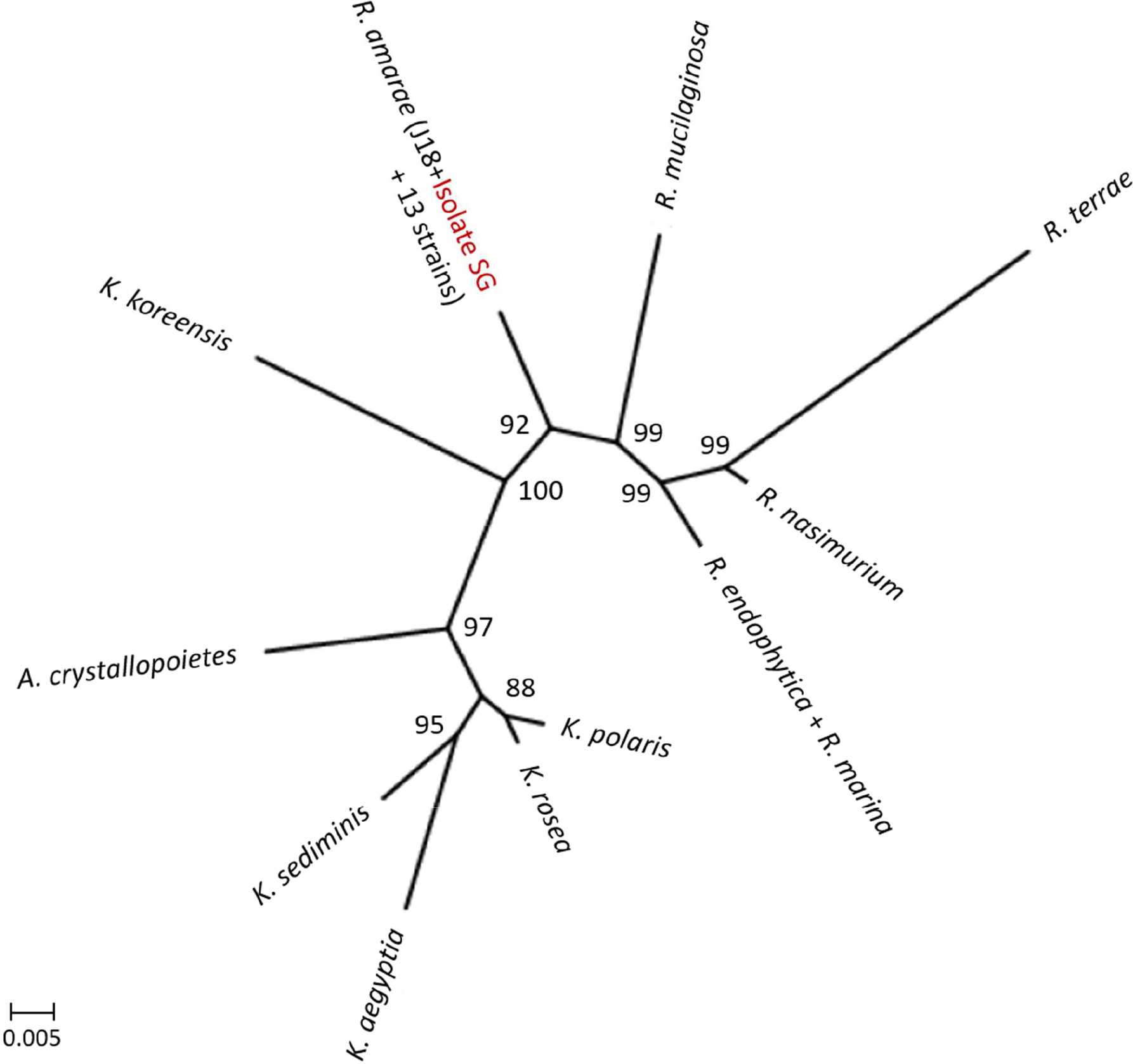
MrBayes phylogram (Phylogeny.fr 2023) for the 16S rRNA gene sequences of the type strains closely related to Isolate SG at identity *I* > 96%. The names of the strains are given in Table 1 and Supplementary Table S1. Numbers are branch support values (Bayesian posterior probability).

**Table 2.** Habitats of *Micrococcaceae* type strains closely related to Isolate SG in the 16S rRNA gene sequence BLAST test (pairwise sequence identity, > 96%)

| No. | Strain | GenBank accession | Length (bp) | Isolation source |
| --- | --- | --- | --- | --- |
| 1 | *R. mucilaginosa* DSM 20746<sup>T</sup> | NR_044873.1 | 1467 | Mucous membranes of the human oral cavity |
| 2 | *R. terrae* L-143<sup>T</sup> | NR_043968.1 | 1436 | Soil in Taiwan |
| 3 | *R. nasimurium* CCUG 35957<sup>T</sup> | NR_025310.1 | 1481 | Mouse |
| 4 | *R. endophytica* YIM 67072<sup>T</sup> | KC806052.1 | 1520 | Aquatic herbaceous plant *Dysophylla stellata* (Lour.) Benth. |
| 5 | *R. marina* JSM 078151<sup>T</sup> | FJ425908.1 | 1443 | Intertidal sediment of the South China Sea |
| 6 | *K. polaris* CMS 76or<sup>T</sup> | NR_028924.1 | 1480 | Antarctic cyanobacterial mat sample |
| 7 | *K. rosea* DSM 20447<sup>T</sup> | NR_044871.1 | 1481 | Found on human skin, in water and in soil |
| 8 | *K. aegyptia* YIM 70003<sup>T</sup> | NR_043511.1 | 1492 | Saline, alkaline desert soil in Egypt |
| 9 | *K. sediminis* FCS-11<sup>T</sup> | NR_118222.1 | 1429 | Marine sediment |
| 10 | *K. koreensis* P31<sup>T</sup> | NR_116745.1 | 1497 | Korean fermented seafood |
| 11 | *A. crystallopoietes* DSM 20117<sup>T</sup> | NR_026189.1 | 1472 | Soil |

## CONCLUSION

The application of phylogenetic taxonomic procedures leads to the improvement of bacterial classification, but even in this case, the need still exists to further clarify the relationships within the taxon we are considering, which encompasses organisms of agricultural, biotechnological, clinical and ecological importance (Nouioui *et al*. 2018).

Phylogenetic studies of our bacterial isolate (Isolate SG) from an *Arabidopsis* suspension culture (Sokolov *et al*. 2021) by using MSAs and comparisons of the SSU rRNA secondary structures have provided convincing evidence for contamination of the published 16S rRNA gene sequence of *R. amarae* type strain J18^T^ (GenBank AY043359.1). Correction of this sequence by deleting the contamination region eliminates contradictions in the obtained array of bioinformatic results. Further, correction of the contaminated sequence yields sequence identity values for the 16S rRNA genes of the isolate and *R. amarae* J18^T^ (and more than a dozen other *R. amarae* members) above the threshold for species demarcation. Thus, the corrected 16S rRNA gene sequence of *R. amarae* type strain J18^T^ should be re-deposited in GenBank for its correct use in various phylogenetic studies.

The phylogram of the 16S rRNA gene sequences of the type strains closely related to *R. amarae* Isolate SG (interspecies sequence identity values, 96–98.7%) united members of the family *Micrococcaceae* (the genera *Rothia, Kocuria* and *Arthrobacter*). A great diversity of habitat conditions was noted for these bacteria, isolated from animals, soil, aqueous media, plant tissues and other sources. This applies, in particular, to *R. amarae* members that are most closely related to the isolate under study by the 16S rRNA gene sequence criterion (sequence identity, ≈100%) assigned to four ecotypes: Antarctic, aquatic, soil and endophytic. The high adaptive potential of bacteria of this species, shown in these observations, enables them to form symbiotic relationships with plants at the endophyte level. *R. amarae* strain SG is likely to be included in their number after additional physiological, biochemical and genetic studies are made.

## ACKNOWLEDGEMENT

We thank Dmitry N. Tychinin (IBPPM RAS) for translating the original manuscript into English.

